# Consumption of Reinforcing Solutions Engages Dynamic Activity of the Prelimbic Cortical Outputs

**DOI:** 10.1101/2023.04.04.535635

**Authors:** Jennifer A. Rinker, Munir Gunes Kutlu, Jason Knapp, Michaela Hoffman, Thomas J. Wukitsch, Erin S. Calipari, Christopher S. McMahan, G. Hamilton Baker, John J. Woodward, Patrick J. Mulholland

## Abstract

The medial prefrontal cortex integrates information about salience and valence of stimuli, including rewarding solutions like alcohol and sucrose, and regulates aspects of alcohol seeking and consumption. However, our understanding of how cortical outputs encode alcohol consumption is limited. Using fiber photometry to measure calcium activity in putative prelimbic (PrL) glutamatergic projection neurons, we show similar but distinct patterns of activity during the peri-consummatory phase in response to consumption of water (non-deprived conditions), ethanol (20% v/v), or sucrose (1% w/v). PrL population activity appears to track hedonic value, as GCaMP6f signals ramped immediately preceding bouts for water, ethanol, and sucrose, and the signal scaled with presumed hedonic value, i.e., water<ethanol<sucrose. Further, using machine learning, population activity of PrL neurons prior to consumption was sufficient to predict both consumption of and distinguish between these different solutions. To assess valence encoding, we adulterated the ethanol solution with quinine, a bitter tastant. In non-dependent mice, calcium activity surrounding drinking bouts was reduced, paralleling the decreased consumption of quinine-adulterated ethanol. This effect was not present in ethanol dependent mice, suggesting altered hedonic value of the adulterated solution either due to reduced sensitivity to the aversiveness of quinine or increased sensitivity to the reinforcing value of ethanol. The global population of PrL glutamatergic neurons also display sustained GCaMP6f “up-states” that last tens to hundreds of seconds, which were longer and larger when consummatory bouts occurred. Overall, our results demonstrate a functional signature in PrL neurons that differs across solutions and is disrupted by ethanol dependence.

## Introduction

The prefrontal cortex (PFC) regulates executive functioning and goal-directed actions[1, 2], and its function can be profoundly impacted by excessive alcohol drinking[3–7]. Alcohol-induced deficits in PFC function can lead to craving and cognitive dysfunction that interfere with recovery efforts in individuals with alcohol use disorder (AUD)[8–10]. With a lifetime AUD prevalence of upwards of 12%[11] and binge drinking rates increasing by 8%[12], understanding alcohol’s impact on cortical function and how the PFC regulates drinking behaviors is critical. The prelimbic (PrL) cortex is a subregion of the medial PFC (mPFC) involved in ethanol seeking behaviors, as well as excessive and compulsive ethanol drinking. Rodent studies have shown that inactivation of the PrL reduces ethanol self-administration in rats[13, 14], and that rats will self-administer ethanol directly into the PrL cortex[15]. Chronic exposure to ethanol in rodents produces functional[16–24], molecular[25, 26], and morphological[4, 19, 23, 24, 27] neuroadaptations in the PrL cortex. Importantly, these adaptations are proposed to be critical drivers of cognitive deficits[24, 28] and compulsive ethanol drinking[7, 26, 29–31].

Neurons in the mPFC can encode anticipatory and goal-directed drinking behaviors[32–38]. A recent study revealed that calcium activity of mPFC glutamatergic neurons that project to the dorsal periaqueductal gray predict future compulsive drinking[39]. Further, a unique pattern of neural activity in mPFC neurons of Wistar rats appears to encode the intention to drink alcohol with increased firing to an ethanol-predictive cue and a peak in neural firing post-consumption[34]. Interestingly, the neural signal encoding the intention to drink alcohol in Wistar rats was blunted in alcohol-preferring P rats[34], and compulsive alcohol drinking was associated with functional changes in PrL neural activity leading to reduced behavioral control[7]. Thus, neural activity in the PrL cortex during active ethanol drinking and adaptations in the PrL produced by chronic ethanol exposure appear to drive ethanol-seeking behaviors.

In the current study, we first sought to compare PrL neural activity patterns surrounding ethanol drinking with the signal for water and sucrose intake using fiber photometry recordings of GCaMP6f calcium transients. Because of the heterogeneous nature of the PrL, we selected fiber photometry to record population activity, as PrL neural populations represent network states[40]. Next, we determined how ethanol dependence alters population activity of PrL glutamatergic projection neurons during voluntary ethanol drinking. Further, to assess changes in valence encoding, we determined how adulterating ethanol with quinine changed the neural activity pattern of PrL neurons before and after the induction of ethanol dependence. Because machine learning (ML) can predict neural activity from behavior[39, 41, 42], we applied two different ML algorithms to determine if the signature of neural activity preceding drinking behavior could predict consumption of different solutions.

## Methods

### Mice

Due to baseline differences in alcohol[43] and sucrose consumption[44, 45] and increased vapor exposure needed to engender dependence-like phenotypes in females[46], these studies used adult male C57BL/6J mice (n = 24; Jackson Laboratories; Bar Harbor, ME; strain # 000664). Mice were group-housed (4/cage) and acclimatized to the colony room for at least one week in a temperature and humidity-controlled AAALAC-approved facility. Mice were maintained on a reverse 12-h light/dark cycle with lights off at 09:00 am and had ad libitum access to food and water. All mice were treated in strict accordance with the NIH Guide for the Care and Use of Laboratory Animals[47] and all procedures were approved by the Medical University of South Carolina’s Institutional Animal Care and Use Committee.

### Stereotaxic Surgery

Mice were deeply anesthetized with vaporized isoflurane (1-3%, SomnoSuite Vaporizer, Kent Scientific) and 200 nl of AAV1-CaMKII-GCaMP6f (Addgene, Catalog #100834-AAV1) was microinfused into the PrL cortex (AP: +1.70; ML: ±0.4; DV: −2.35; in mm relative to bregma) at a rate of 100 nL/min. The microinjection needle remained in place for 5 min post-infusion. After infusion, optical ferrules (400 *μ*m, 0.48 NA, 2.5 mm OD ceramic; Doric Lenses) were implanted above the infusion site in one hemisphere and secured via Herculite dental resin (Kerr Dental). Mice were transferred to individual home cages and allowed to recover for 4-6 weeks. Mice were habituated to handling, tethering to patch cords and lickometer circuitry (MedAssociates) during the recovery period.

### Ethanol drinking and CIE Exposure

Binge-like ethanol consumption was induced using a modified 4-d drinking-in-the-dark (DID) protocol[48–51]. Fiber photometry recordings occurred on the 4^th^ day of each cycle of DID, and the drinking sessions lasted for a 2-hr period beginning 3 hr into the dark cycle. In the first cohort of mice (n = 9 mice), standard water bottles were replaced with a single bottle of water for the test drinking sessions in week 1. For test weeks 2 through 4, water was replaced with a single bottle containing 20% ethanol (v/v). After a 2- to 4-wk abstinence period from ethanol drinking, mice were given access to a single bottle containing 1% sucrose (w/v) for drinking sessions. In the second cohort (n = 10 mice), mice were allowed to drink ethanol in the DID protocol for 3 weeks prior to and 1 week after exposure to chronic intermittent ethanol (CIE) in vapor inhalation chambers. Mice in the 3^rd^ cohort (n = 5 mice) were treated in the same manner with the addition of a test drinking session where 250 *μ*M quinine was added to the ethanol solution before and after CIE exposure. Air or ethanol vapor was delivered 16 hr/day for four consecutive days followed by three days of abstinence before beginning the next cycle of exposure, which was repeated for 4 weeks[52–54]. Before entry into chambers, CIE mice received a priming dose of ethanol to initiate intoxication (1.6 g/kg; 8% w/v in sterile saline; 20 ml/kg injection volume; IP) with pyrazole (1 mmol/kg) to slow alcohol metabolism. Air-exposed mice were handled similarly but received pyrazole in sterile saline. Mice were exposed to ethanol vapor at concentrations that yield blood ethanol concentrations (BECs) in the range of 175-225 mg/dL. BECs in samples taken from the retroorbital sinus were determined using an alcohol analyzer (Analox Instruments, Stourbridge, UK)[52].

### Fiber photometry recordings

GCaMP6f transients were recorded during 2-hr drinking sessions using a fiber photometry system consisting of 405 and 470 nm light-emitting diodes (LED) coupled into a 400 *μ*m 0.48 NA optical fiber (Doric Lenses) connected to an integrated fluorescent mini-cube (Doric Lenses). Excitation power was set to 15 *μ*W for each channel. The optical fiber from the mini-cube connected to a pigtailed rotary joint (Doric Lenses) and the optic fiber from the rotary joint connected to the implanted fiber using a ceramic sleeve or pinch connector (Thorlabs). The rotary joint was suspended in a custom-build balance arm that allowed free movement. Emission light was focused onto a photodetector (Newport model 2151; DC low setting) low-passed filtered at 3 Hz and sampled at 6.1 kHz by a RZ5P lock-in digital processor (TDT) controlled by Synapse software (TDT). Excitation light was sinusoidally modulated at 531 Hz (405 nm) and 211 Hz (470 nm) via software control of an LED light driver (Thorlabs). Real-time demodulated emission signals from the two channels were acquired at a frequency of 1017.25 Hz and stored offline for analysis. Data from lickometers were also collected via TTL inputs to the digital processor. All patchcords were photobleached using a 405 nm LED at 500 μW power for >12 h every 3-4 days to reduce autofluorescence. Raw photometry data were processed and time-locked to TTLs using custom written functions in MATLAB (Mathworks) software. Because the 405 nm and 470 nm signals showed differences in photobleaching over time (slope, mean ± SEM; 405 nm −0.056 ± 0.001, 470 nm −0.0042 ± 0.0004, paired t-test, t_44_ = 2.347, p = 0.023, n = 45 recordings), the signals for each channel were first fitted to a polynomial versus time and then subtracted from one another to calculate the %Δ*F*/*F* time series[55]. Peaks in GCaMP6f fluctuations where the median absolute deviation (MAD) was >2 were quantified using the ‘findpeaks’ function[56–58]. We determined the peak in the normalized GCaMP6f signal that occurred 3 seconds prior to and after each licking bout. The minimum GCaMP6f value was determined during each drinking bout to quantify the drop in the signal that occurred during active licking. Finally, the average GCaMP6f signal preceding each licking drinking bout was determined in 10 sec epochs for 1 min. The area under the curve (AUC) for the GCaMP6f signal was calculated for the 30 sec period preceding and following each drinking bout. A licking ‘bout’ was identified if lick frequency was >4.5 Hz, the interlick interval was < 500 ms, and the bout duration was > 500 ms[59–61]. Prolonged GCaMP6f up-states were defined as increases in the signal above one MAD that lasted >1 second. Slope and intercept variables for the 30 seconds preceding each drinking bout were derived from a linear mixed-effects model (“lme4” package[62] and optimized using the “nlminb” method; “optimx” package[63] in R).

### Machine Learning

A machine learning classifier support vector machine (SVM[41]) was used to determine if the GCaMP6f signal 30 seconds prior to a drinking bout could predict consumption of water, ethanol, or sucrose. A custom MATLAB code was used to create randomly selected training and test data sets for SVM analysis using the best predictive features in a kernel function (radial basis function, RBF) to find the optimal hyperplane between binary prediction options[41]. Then, the trained model was used to predict the accuracy of classifying the test data set across 20 trials. We first compared pre-bout data for each drinking solution with random 30-second epochs using the same number of random epochs as drinking bouts within each recording. Next, we analyzed whether four features of the 30 second pre-bout data (slope, intercept, peak 3 seconds prior to a bout, and AUC) predicted drinking for water, ethanol, or sucrose. Data for the features were log2-transformed, with the exception of slope given the presence of negative values for some water drinking bouts. All data sets were also analyzed using randomly scrambled group classifications.

For multi-class classification, an extreme gradient boosting algorithm (i.e., XGBoost) was selected because of its ability to work with small, sparse data sets and more accurately predict neural data compared with other approaches, included neural networks and generalized linear models[42, 64]. XGBoost also allowed us to determine when important features occurred while also comparing across all three solutions. We first reduced the dimension of the feature space via functional data analysis techniques as outlined in[65] by viewing the 30 second GCaMP6f signal as data that is arising from an unknown function. Next, an estimator of this function was developed via a B-spline approximation using 300 basis functions. The coefficients from these approximations were then used as the feature set, which reduced the feature space from >30,000 variables to 300, while preserving virtually all data available. Custom R code was used to create randomly selected training and test data sets. Model training made use of repeated 10-fold cross validation, repeated 10 times, to tune the number of regression trees and tree depth of the XGBoost model. To evaluate prediction accuracy, the trained model was used to predict the accuracy of classifying the test data for 20 trials. This entire process was repeated 100 times. Variable importance was assessed via VIP scores for each model. To aggregate the VIP scores, we computed the average VIP score for each feature and then normalize these scores to exist between 0 and 1 by dividing by the max.

### Histology

GCaMP6f expression and location of optical ferrules were identified using histological techniques. Mice were transcardially perfused with Dulbecco’s PBS and 4% paraformaldehyde and ferrule placements were verified via the visualization of optical fiber tracks and viral vector transduction efficiency via native GCaMP6f expression. Tissue sliced at 40*μ*m on a vibratome (Leica VT 1000S, Leica Biosystems), and images were acquired using a Zeiss LSM 880 confocal microscope.

### Statistics

All behavioral and photometry data were analyzed as a general linear mixed model using Tukey post-hoc tests, when applicable, following our previous methods[66, 67]. For all repeated measures designs, data were nested within mouse and time. Data are reported as mean ± SEM, and statistical significance was established with α = 0.05.

## Results

### Voluntary drinking

Mice that were allowed to drink water, 20% ethanol, or 1% sucrose in their home cages consumed more sucrose (in ml/kg) than water and ethanol (F_2,14_ = 4.52, p = 0.031; **Fig 1A**). There were significant differences in bout duration (**Fig 1B**; F_2,13_ = 39.45, p < 0.0001) and frequency of licking behavior (**Fig 1C**; F_2,13_ = 5.99, p = 0.014) for the three solutions. Posthoc comparisons showed that bout duration was longer for ethanol than sucrose and water (**Fig 1B**) and bout duration for sucrose was longer than water (**Fig 1B**), but lick frequency for ethanol was slower than that for water or sucrose (**Fig 1C**).

**Figure 1.**
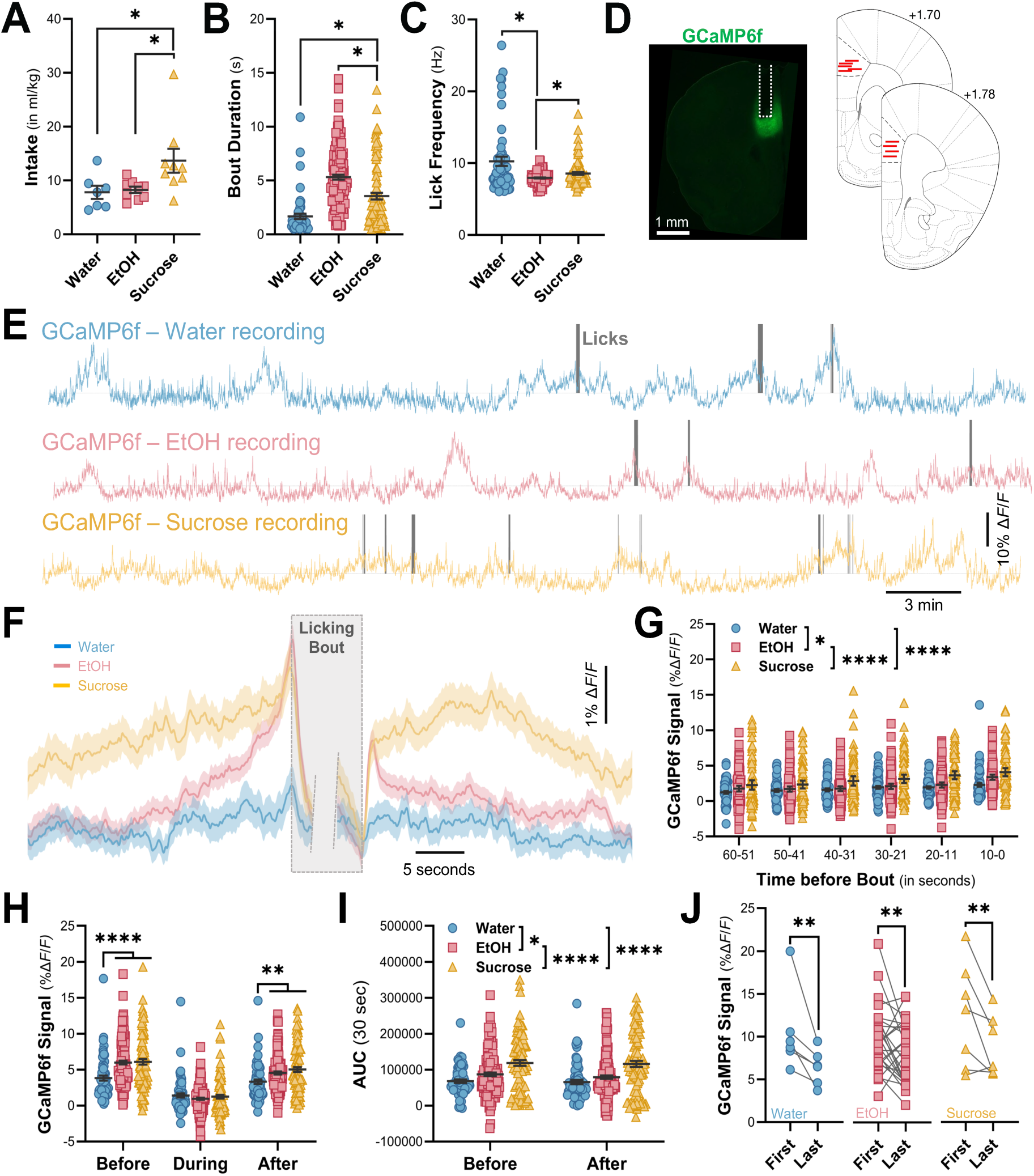
PrL cortex GCaMP6f signaling during water, ethanol (EtOH), and sucrose drinking. ***A-C***, Intake, bout duration, and lick frequency for water, ethanol (20%, v/v), and sucrose (1%, w/v) drinking solutions. ***D***, Representative image of GCaMP6f expression in PrL cortex and location of the chronic optical ferrule implant. ***E***, Representative recordings of normalized GCaMP6f during drinking sessions for water, EtOH, and sucrose. Licks on the sipper tube are demarked by gray vertical lines. ***F***, Averaged GCaMP6f signal prior to and following licking bouts for the three drinking solutions. Averaged traces show mean (solid line) with standard error of the mean (shading). ***G***, Binned GCaMP6f signal preceding licking bouts. Features of the GCaMP6f signal surrounding licking bouts for the different drinking solutions. ***H***, Peak in the GCaMP6f signal 3 sec before and after licking bouts, and the minimum signal during drinking. ***I***, Area under the curve (AUC) before and after licking bouts. ***J***, Comparison of the peak signal preceding the first and last drinking bout for each solution. \**p* < 0.05; \*\**p* < 0.01; \*\*\**p* < 0.001; \*\*\*\**p* < 0.0001.

### Prelimbic GCaMP6f signaling surrounding drinking bouts

Representative traces of the GCaMP6f signal and licking bouts for water, ethanol, and sucrose drinking sessions are shown in **Fig 1E**. Consistent with evidence of increased neural activity during goal-directed behaviors[68], we observed an increase in the PrL GCaMP6f population signal preceding drinking bouts (**Fig 1F**). When analyzing the averaged signal in 10-sec epochs for a 1-min period before each bout, there was a significant main effect of time (F_5,45_ = 15.40, p < 0.0001) and drinking solution (F_2,13_ 41.00, p < 0.0001). Posthoc comparisons revealed that the signal for sucrose was significantly higher than ethanol and water (p < 0.0001), and the signal for ethanol was significantly higher than water (p = 0.013; **Fig 1G**). The signal in the 10-sec epoch preceding a bout was significantly higher than the other epochs (p < 0.01), and the 20-11 and 30-21 second epochs were significantly higher than the epochs between 60 and 41 seconds (p < 0.05).

Because the signal was significantly elevated for both ethanol and sucrose immediately preceding and following a bout, we analyzed the signal before, during, and after drinking bouts, which revealed a significant solution by time interaction (F_4,26_ = 5.69, p = 0.002; **Fig 1H**). For water only bouts, posthoc comparisons showed that the signal during the bout was significantly lower than the signal before and after the bout (p < 0.01). For ethanol and sucrose bouts, the signal was significantly different from each of the other time points (before>after>during; p < 0.05). Posthoc comparison across solutions showed that the signal before an ethanol or sucrose bout was significantly increased compared with the signal before and after a water bout (p < 0.0001 and p < 0.01, respectively). When considering the AUC of the signal 30 seconds prior to and after a licking bout, there was a significant difference across the 3 solutions (F_2,13_ = 27.50, p < 0.0001, **Fig 1I**). Post-hoc analyses revealed that the AUC for ethanol was higher than water (p = 0.034) and the AUC for sucrose was higher than both ethanol and water (p < 0.0001). Because reward signaling for sucrose drinking tracks with satiation[58], we compared the peak signal preceding the first and last bout that occurred within each drinking session across all three solutions. There was a significant main effect of bout (F_1,9_ = 12.22, p = 0.007), and consistent with previous evidence, the peak preceding the first bout was larger than the peak for the last bout (**Fig 1J**).

Since mice consumed nearly twice as much sucrose as ethanol or water, one potential concern is that the elevated signal preceding a sucrose bout is inflated by ramping of activity from overlapping licking bouts. To address this, we extracted and analyzed the signal for licking bouts without overlap. The averaged signal for non-overlapping licking bouts for water, ethanol, and sucrose is shown in **Fig 2A**. When analyzing the ramping activity that preceded non-overlapping bouts (**Fig 2B**), there was a significant main effect of solution (F_2,525.5_ = 23.10, p < 0.0001) and time (F_2,537.2_ = 26.31, p < 0.0001). Posthoc analysis showed that sucrose was significantly different from ethanol and water (p < 0.0001) and that ethanol was significantly different from water (p < 0.001). The signal in the 10-sec epoch preceding non-overlapping bouts was significantly higher than the other epochs (p < 0.0001). When considering the signal surrounding drinking bouts, there was a significant solution by time interaction (F_4,539.38_ = 5.493, p < 0.001; **Fig 2C**). For the signal prior to the bout, posthoc analysis revealed that the signal for sucrose was higher than ethanol (p = 0.008) and water (p < 0.0001) and that ethanol was higher than water (p < 0.05).

**Figure 2.**
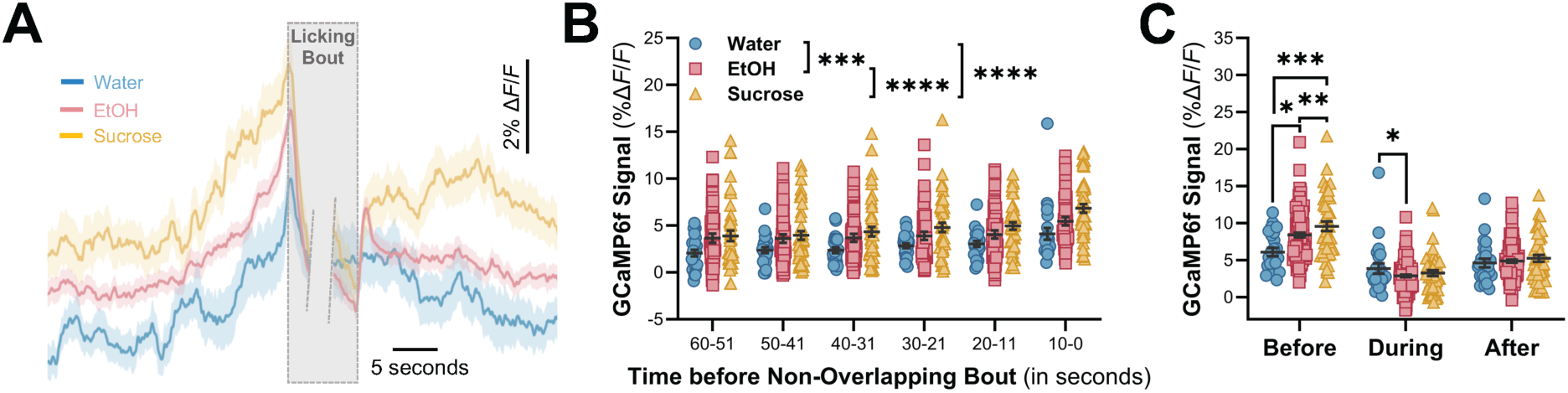
GCaMP6f signaling in the PrL cortex during non-overlapping licking bouts for water, ethanol (EtOH), and sucrose. ***A***, Averaged GCaMP6f signal prior to and following non-overlapping licking bouts for the three solutions. Averaged traces show mean (solid line) with standard error of the mean (shading). ***B***, GCaMP6f signal preceding non-overlapping licking bouts in 10 second epochs. ***C***, Non-overlapping GCaMP6f signal before, during, and after non-overlapping drinking bouts. \**p* < 0.05; \*\**p* < 0.01; \*\*\**p* < 0.001; \*\*\*\**p* < 0.0001.

### Machine learning and functional signature in PrL projection neurons

These results suggest that the GCaMP6f signal, particularly the 30 seconds prior to the start of drinking bouts, may distinctly encode the three different solutions. To test this, we first applied a SVM classifier to determine whether the signal could predict drinking for the individual solutions. The SVM predicted when mice were drinking water (74% accuracy; t_99_ = 17.70, p < 0.0001), ethanol (77.7% accuracy; t_99_ = 19.22, p < 0.0001), and sucrose (70.8% accuracy; t_99_ = 16.00, p < 0.0001) compared with random epochs extracted from their respective drinking sessions (**Fig 3A**). Using XGBoost, we were able to distinguish between the 30 seconds of pre-bout data for three solutions and random epochs when analyzed simultaneously. XGBoost identified 20 time points of high importance factors in the 30 seconds of data preceding a drinking bout (**Fig 3B**). A variable importance plot shows that most of these factors reside within 10 seconds prior to initiation of licking behavior, and the factors with the highest importance reside within ∼5 seconds of bout initiation (**Fig 3B**). XGBoost analysis predicting the three drinking solution and random data from one another demonstrated modest, but highly significant accuracy (i.e., 62.4%) for (t_198_ = 14.83, p < 0.0001; **Fig 3C**).

**Figure 3.**
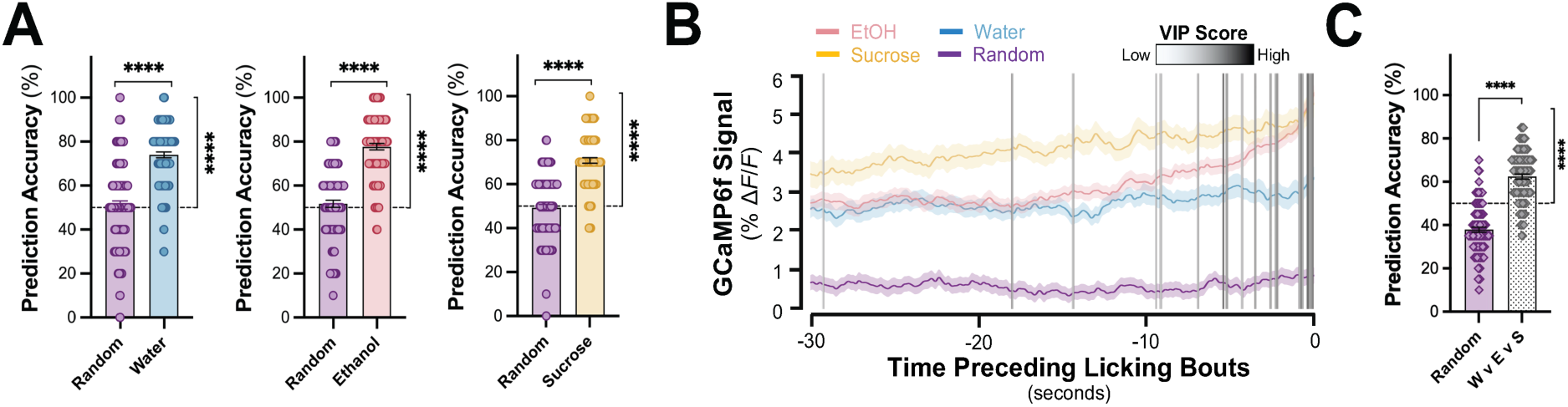
Prediction of different drinking solutions using machine learning approaches. A, Prediction accuracy of the support vector machine for the three different solutions vs. random condition. Accuracy for scrambled (Scr) comparison is also shown. B, Variable importance plot (VIP) locations from the XGBoost algorithm. C, Prediction accuracy for the three different solutions and the random condition. ****p < 0.0001.

### Calcium event dynamics and prolonged GCaMP6f up-states

Next, we determined if ethanol or sucrose drinking altered peaks in GCaMP6f fluctuations (i.e., ‘events’) in the PrL cortex (**Fig 4A,B**). The frequency (F_2,16_ = 0.63, p = 0.54), amplitude (F_2,16_ = 1.56, p = 0.24), and width (F_2,16_ = 0.70, p = 0.51) of GCaMP6f events during ethanol and sucrose drinking sessions were not significantly different from water drinking sessions (**Fig 4C-E**). In addition to peaks in GCaMP6f fluctuations, we observed sustained increases in the signal that were reminiscent of persistent cortical up-states[69], but lasted tens to hundreds of seconds. As suggested by the representative traces shown in **Fig 1E**, licking bouts for water, sucrose, and ethanol appeared to occur during prolonged up-states. The mean signal, duration, and AUC of prolonged up-states that did and did not contain licking bouts differed dramatically. For each of these measures, there was a significant main effect revealing increases when bouts were present (*mean*: F_1,9_ = 58.5, p < 0.0001; *duration*: F_1,9_ = 74.88 p < 0.0001; *AUC*, F_1,9_ = 45.07, p < 0.0001; **Fig 4F-H**). There was also a significant main effect of solution (F_2,17_ = 7.47, p = 0.005) for the signal during prolonged up-states, with posthoc analyses indicating that the up-state mean for ethanol and sucrose were higher than water (p < 0.05; **Fig 4F**).

**Figure 4.**
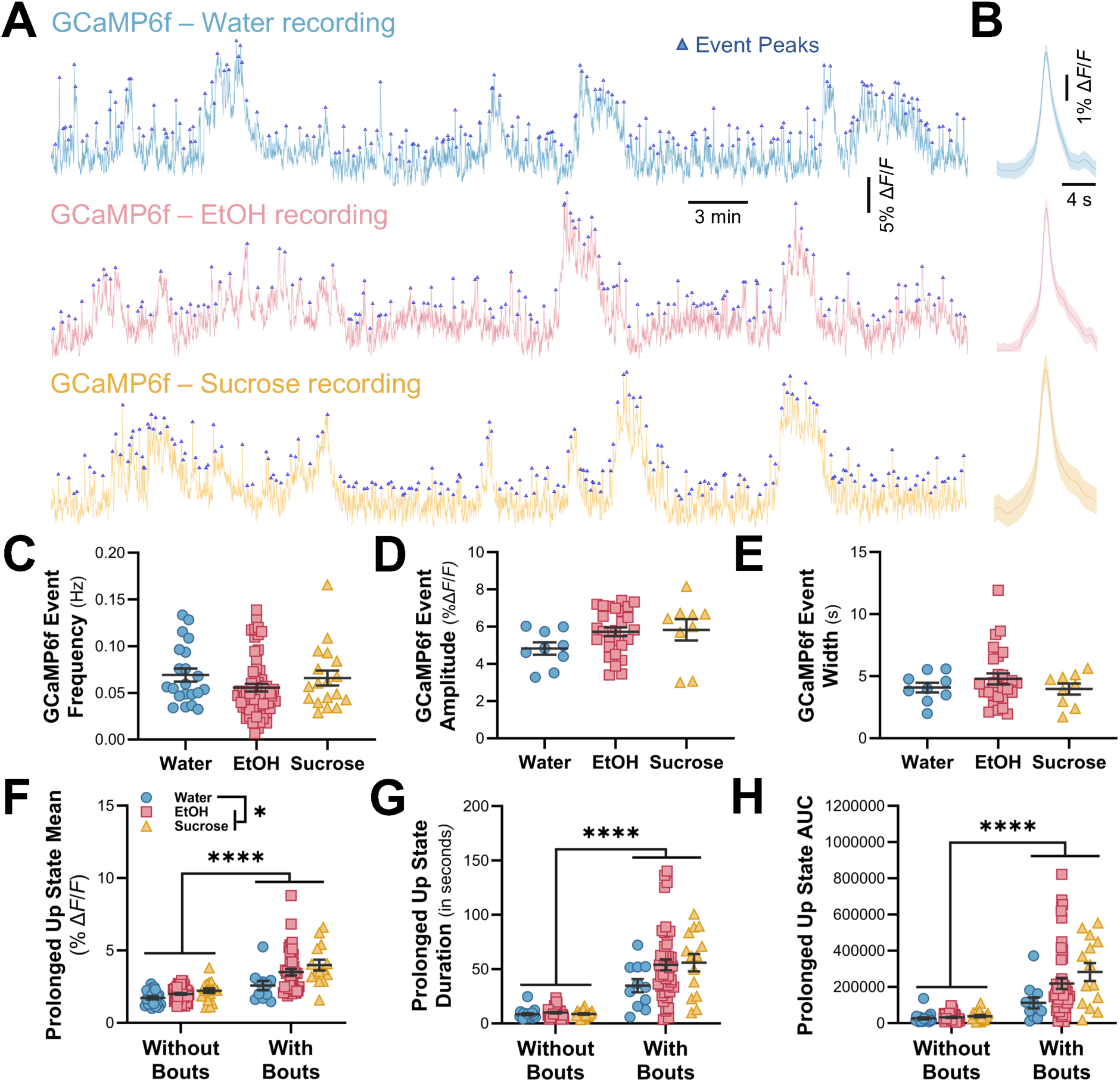
GCaMP6f events are similar across drinking solutions. ***A-B***, Representative traces showing GCaMP6f events and averaged events for water, EtOH, and sucrose. Averaged events show mean (solid line) with standard error of the mean (shading). ***C-E***, GCaMP6f event frequency, amplitude, and width were similar. Features of prolonged GCaMP6f up-states differed with and without drinking bouts. ***F-H***, Mean, duration, and AUC of prolonged up-states were greater when it contained a licking bout. Prolonged up-state mean was also greater for EtOH and sucrose vs. water. \**p* < 0.05; \*\*\*\**p* < 0.0001.

### CIE exposure, aversion-resistant drinking and prelimbic GCaMP6f signaling

Given the distinct PrL calcium signal in mice drinking ethanol, we next wanted to determine how ethanol dependence alters the signal surrounding drinking bouts. While ethanol drinking was similar during baseline, CIE exposure increased drinking in ethanol dependent mice during the post-vapor test drinking sessions (treatment x time interaction: F_1,8_ = 19.12, p = 0.002; n = 5 mice/group; **Fig 5A**). Consistent with our initial recordings, we observed peaks in the signal prior to and after drinking bouts for ethanol prior to CIE (**Fig 5B**). The signal surrounding licking bouts pre- and post-Air or -CIE was analyzed by a repeated-measures 3-way mixed linear model. There was a significant time (pre- and post-vapor) x treatment (air vs CIE) x signal (before, during, after licking) 3-way interaction (F_2,575_ = 3.06, p = 0.047, n = 5 mice/group). Analyses of simple interactions before, during, and after licking bouts revealed a significant interaction between treatment and time for the signal before (F_1,607_ = 7.45, p = 0.007), but not during (F_1,607_ = 0.14, p = 0.713) or after (F_1,607_ = 3.19, p = 0.075) drinking bouts. The second-order simple interaction analysis demonstrated that the signal prior to licking bouts significantly decreased (p = 0.0005) in the air exposed controls from baseline, whereas CIE exposed mice showed no change from baseline (p = 0.901). For the signal during drinking bouts, there was a significant main effect of time (F_1,607_ = 9.92, p = 0.002), but not treatment (F_1,607_ = 0.03, p = 0.868). The post-CIE signal during the bout in both groups was significantly lower than the baseline values. There were also main effects of time (F_1,607_ = 12.05, p = 0.0006) and treatment (F_1,607_ = 9.72, p = 0.0019) for the signal that occurred after a drinking bout showing a reduction in the amplitude compared with the pre-vapor exposure values and the CIE-exposed mice, respectively. These data are represented in **Fig 5E** as the raw change in the signal (post-vapor minus baseline) in the two treatment groups. Closer inspection of the signal in **Fig 5D** suggested that the change in signal from peak before to valley during consumption was greater in CIE-exposed mice. To test this directly, we calculated the difference in the signal before minus during licking bouts, and the two-way, repeated-measures analysis revealed a significant interaction between time and treatment (F_1,8_ = 7.67, p = 0.024). Post-hoc analyses indicated that the difference in the signal drop was significantly amplified in the CIE exposed mice (p = 0.039, **Fig 5F**), but not the air vapor exposed controls (p = 0.142).

**Figure 5.**
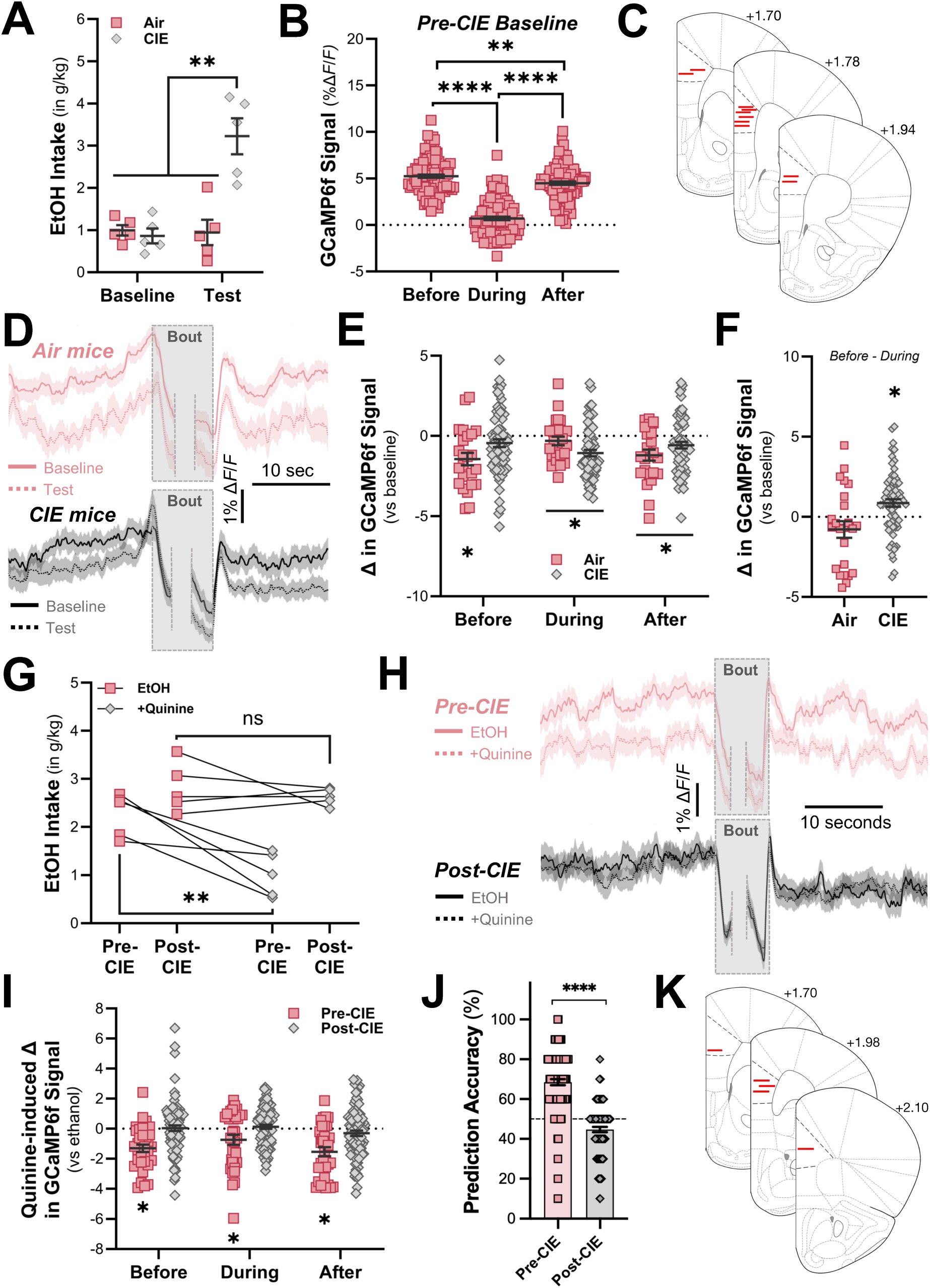
Effects of CIE on PrL GCaMP6f activity. ***A***, EtOH intake in Air and CIE exposed mice at baseline and during the post-CIE test. ***B***, Quantified GCaMP6f signal before, during, and after bouts of licking for EtOH prior to CIE exposure. ***C***, Schematic of location of the chronic optical ferrule implants. ***D***, Averaged GCaMP6f signal surrounding licking bouts for EtOH in Air (red) and CIE (gray) before and after CIE (or Air) exposure. ***E***, Change in GCaMP6f signal from baseline. ***F***, Difference from peak before to min during bouts of licking for EtOH. \**p* < 0.05, \*\**p* < 0.01, \*\*\*\**p* < 0.0001. EtOH dependence blocks quinine-induced PrL GCaMP6f signal reduction. ***G***, EtOH intake pre-CIE and post-CIE (red), and during quinine-adulteration test (gray). ***H***, Averaged GCaMP6f signal for EtOH and quinine-adulterated EtOH bouts pre- and post-CIE. ***I***, Quinine-induced change in GCaMP6f signal before, during, and after licking bouts pre- and post-CIE. ***J***, SVM prediction accuracy for distinguishing EtOH bouts from EtOH+quinine bouts pre- and post-CIE. ***K***, Schematic of location of the chronic optical ferrule implants. \**p* < 0.05, \*\**p* < 0.01, \*\*\*\**p* < 0.0001.

Because the mPFC controls excessive and compulsive-like ethanol drinking[7, 39, 70], we next determined how quinine adulteration of ethanol affected the PrL signal in ethanol dependent mice. Prior to CIE exposure, the addition of 250 *μ*M quinine to the ethanol drinking solution significantly reduced intake in mice (interaction: F_1,4_ = 8.03, p = 0.047; post-hoc, p = 0.009; n = 5 mice; **Fig. 5G**). The signal surrounding licking bouts for ethanol and quinine-adulterated ethanol pre- and post-CIE exposure were analyzed using a within-subjects, three-way mixed linear model. The 3-way interaction was not significant (F_2,713_ = 0.50, p = 0.609), but there was a significant treatment x drinking solution interaction (F_1,713_ = 10.63, p = 0.001). Post-hoc analysis from the simple effects analysis demonstrated that the signal was significantly reduced when mice consumed ethanol adulterated with quinine during the pre-CIE drinking test (p = 0.009, **Fig 5I**). In contrast, these mice consumed equal amounts of ethanol when quinine was added to the drinking solution during the post-CIE test (posthoc, p = 0.531; **Fig 5G**), and the signal surrounding drinking bouts was unaltered (p = 0.748, **Fig 5I**). The SVM predicted the quinine-induced change in the calcium signal prior to, but not after CIE exposure (t_198_ = 11.59, p < 0.0001; **Fig 5J**).

## Discussion

In this study, we provide experimental and computational evidence that PrL pyramidal neuron population activity tracks consumption of different rewarding solutions. Strikingly, ramping of PrL activity predicts drinking behavior for three solutions with differing perceived rewarding properties. Moreover, we showed that mice predominantly consumed fluid when the PrL population signal was in a prolonged upstate, and features of the pre-bout ramping signal could accurately discriminate between the three drinking solutions. Consistent with declining VTA signaling across sucrose licking within a drinking session reflecting satiety-induced reductions in reinforcing value[58], we observed a decrease in the peak calcium signal for all solutions across time. While photometry signal can decline during extended recording sessions and we cannot exclude the possibility that a component of this reduction reflects photobleaching of the calcium sensor rather than solely within-session satiety, it should be noted that the amplitude of other calcium events was sustained across the entire recording session. Another main finding from our study is that the pre-bout PrL signal parallels escalated and compulsive drinking behavior in a mouse model for the study of AUD, i.e., remains elevated in dependent mice that show aversion-resistant (quinine adulteration) consumption of ethanol, but is diminished in control mice sensitive to quinine adulteration. Overall, our results provide evidence for a functional signature in the PrL cortex that tracks solution-specific engagement, perhaps contributing to the intention to drink, especially when mice are rendered dependent upon ethanol.

### Anticipatory PrL signaling and licking behavior

Our data extend previous findings by providing evidence for anticipatory signaling in the population of PrL glutamatergic projection neurons in non-deprived, freely moving mice. The population calcium signal ramped prior to drinking bouts, peaking just prior to bout initiation for ethanol or sucrose. The ramping of the cortical GCaMP6f signal prior to goal-directed behavior is consistent with single unit data[68, 71–73], although our photometry data showed longer ramping timescales preceding consumption, perhaps reflecting anticipatory signaling or motor planning at the population level. Additionally, because licking dynamics differed across solutions, it is also possible that components of the pre-bout ramping signal reflect motor preparation or sensorimotor processes associated with the initiation of licking behavior rather than purely anticipatory reward processing. Future work combining photometry with high-resolution behavioral tracking or motor controls will be important to further dissociate these contributions.

Additionally, we also observed ‘prolonged’ up-states in the GCaMP6f signal over the course of the drinking sessions that last longer (>60 seconds) than traditionally defined ‘persistent’ cortical up-states[69]. Interestingly, drinking bouts frequently occurred during these prolonged up-states, and characteristics of the prolonged up-states in the PrL population were quantitatively different in the presence of a drinking bout, especially for ethanol and sucrose. While the prolonged up-states appear important for signaling drinking, some prolonged up-states occur in the absence of drinking, suggesting that additional signaling outside of the PrL or within a specific PrL projection is necessary. Together, these data suggest that population calcium activity in PrL glutamatergic neurons is associated with the initiation of drinking behavior and behavioral engagement with reinforcing solutions, including necessary solutions, such as water under non-deprived conditions, and solutions with rewarding properties. It is also important to note that the voluntary consumption tests were conducted sequentially (water, ethanol, then sucrose), and therefore differences in PrL signaling across solutions may partially reflect experience-dependent changes in learning or task familiarity over time. While these data demonstrate clear solution-specific differences in neural activity patterns, future studies employing counterbalanced presentations will be required to fully isolate solution identity from potential contextual or temporal learning effects.

### PrL neurons encode hedonic value

Consistent with evidence that single unit activity in the mPFC encodes the magnitude of the reward[36–38], the anticipatory photometry signal preceding drinking bouts appeared to track the perceived hedonic value of each solution tested in this study. We found that multiple features (pre-bout ramping, AUC, peak signal) of the GCaMP6f signal differed across the three drinking solutions. Whereas peaks for ethanol and sucrose were similarly increased above peaks surrounding water bouts, the other measures showed differences across all three solutions. Importantly, we interpret these differences cautiously. Although ethanol and sucrose are commonly considered rewarding in rodent paradigms, the present experiments did not directly measure hedonic evaluation of taste preference. Therefore, the neural differences observed here likely reflect solution identity or relative reinforcing value rather than a direct encoding of hedonic value per se.

Presumably, ethanol is more rewarding than water, reflected by longer bout duration compared to non-water deprived conditions, and these same mice consumed more sucrose (in ml/kg) than ethanol and water. To further support evidence for solution-specific encoding by PrL neurons, machine learning analyses predicted drinking bouts against both random epochs of the calcium signal and each of the three solutions. In addition to these computational findings, pairing ethanol with the aversive, bitter tastant quinine in non-dependent mice not only produced the expected reduction in ethanol consumption, but also reduced the GCaMP6f signal surrounding drinking bouts. These results suggest that PrL population activity is sensitive to changes in motivational state or sensory context associated with the drinking solution.

Finally, the peak GCaMP6f signal preceding drinking declined within individual drinking sessions for all solutions, which may reflect declining motivational state, sensory adaptation, or technical factors such as photobleaching, and may explain the modest, but still highly significant, differences identified by the ML analyses. Thus, these studies provide multiple lines of evidence that population activity of PrL projection neurons differentiates between drinking solutions and behavioral states associated with their consumption.

### CIE exposure and anticipatory PrL signaling

Previously, calcium activity in PrL neurons projecting to the dorsal periaqueductal gray predicted future compulsive ethanol drinking[39]. While we did not design our study to predict subsequent drinking phenotypes, we measured population activity of PrL neurons following CIE exposure when mice display a compulsive-like phenotype[74–77]. In the air control mice, the GCaMP6f signal surrounding ethanol drinking bouts decreased across time, which is consistent with previous evidence of signal habituation to repeated reward access[58]. As noted above, this reduction may reflect motivational changes, neural adaptations, or photometry signal decay over prolonged recordings, and these possibilities should be interpreted prudently.

Whereas quinine adulteration reduced drinking and the calcium signal in control mice, the addition of quinine did not affect drinking in CIE-exposed mice, and the calcium signal surrounding drinking bouts remained elevated, possibly reflecting known PrL adaptations produced by CIE exposure[4, 18, 24, 78]. Notably, CIE did not affect other measures of the normalized GCaMP6f signal or the raw values, suggesting that the CIE-induced change in signaling may reflect altered motivational processing or reduced sensitivity to aversive feedback associated with ethanol consumption in PrL neurons. These effects of CIE on signaling surrounding ethanol drinking bouts appear specific to the PrL, at least when compared with the lateral orbitofrontal cortex, which did not reveal a change in the anticipatory signal in ethanol dependent mice[74]. However, unlike the PrL cortex, CIE exposure increased frequency of GCaMP6f events in the OFC[74], demonstrating selective adaptations in frontal cortex after induction of ethanol dependence.

While our pre-bout data are consistent with an ‘intention to drink’ signal in the mPFC of outbred Wistar rats[34], the intention signal after CIE markedly differs from that of alcohol-preferring P rats selectively bred for high ethanol drinking that are commonly used to study familial risk associated with AUD[79]. The intention signal preceding ethanol drinking is blunted in P rats compared with Wistar rats and the GCaMP6f signal in our study remains elevated in CIE exposed mice compared with air controls. These data suggest that genetic predisposition and induction of ethanol dependence may exert differential control over excessive drinking through divergent signaling mechanisms in the PrL cortex. Although there are major differences between these two studies that could explain the differences in P rats and CIE-exposed mice (e.g., species, recording technique, and ethanol paradigm), we would argue that both studies provide evidence that aberrant signaling within the PrL cortex leads to excessive ethanol intake. Alternatively, it is possible that reorganization of other neural circuits or regions innervated by the PrL would show consistent neural firing patterns in CIE exposed mice and P rats that could explain the excessive drinking phenotype. Our findings demonstrating an impaired intention signaling and altered response to quinine adulteration in the PrL cortex are also complementary to a recent study showing the strengthening of neural representation of ethanol seeking in the mPFC of compulsive drinking rats when challenged with quinine-adulterated ethanol[7].

In summary, monitoring population calcium activity in the PrL cortex identified a pre-bout signal that varied with different solutions and tracked changes in compulsive-like ethanol drinking in dependent mice. Multiple ML algorithms supported solution-specific patterns of neural activity in PrL projection neurons associated with drinking. Finally, we identified an aberrant ‘intention to drink’ signal in the PrL cortex that could serve as a mechanism driving excessive and compulsive-like ethanol drinking in dependence, while acknowledging that future studies will be required to disentangle contributions of motor preparation, contextual learning, and motivational state to the observed neural dynamics.

## Conflict of Interests

The authors have nothing to disclose.

## Author Contributions

Conceptualization, JAR, PJM; Investigation, JAR, JK; Data analysis, JAR, MGK, MH, TJW, ESC, CSM, GHB, PJM; Writing, JAR, MGK, CSM, PJM; Editing, JAR, MGK, ESC, CSM, GHB, JJW, PJM; Funding, JAR, MGK, & PJM.

## Funding

These studies were supported by the Charleston Alcohol Research Center (AA010761), the INIAstress Consortium - AA020930 (PJM) and NIH Grants: OD021532 (PJM), AA023288 (PJM), AA025110 (JAR), and MH132052 (MGK).

## Notes

### Competing Interest Statement

The authors have declared no competing interest.

### Summary of Updates

In the revised manuscript, we provide additional analyses resulting in slight changes in the interpretation of the findings.

